# Effects of *Platycodon grandiflorum* on Gut Microbiota and Immune System of Immunosuppressed Mouse

**DOI:** 10.1101/2021.05.07.443070

**Authors:** So Yun Jhang, Sung-Hyen Lee, Eun-Byeol Lee, Ji-Hye Choi, Sohyun Bang, Misun Jeong, Hwan-Hee Jang, Hyun-Ju Kim, Hae-Jeung Lee, Hyun Cheol Jeong, Sung Jin Lee

**Affiliations:** Interdisciplinary Program in Bioinformatics, Seoul National University, Kwanak-Gu, Seoul, Korea; National Institute of Agricultural Sciences, Rural Development Administration, Iseo-myeon, Wanju-Gun, Jeollabuk-do, Korea; Institute of Bioinformatics, University of Georgia, Athens, GA, USA; eGenome, Seoul, Korea; World Institute of Kimchi, Nam-Gu, Gwangju, Korea; Department of Food and Nutrition, Gachon University, Seongnam, Gyeonggi, Korea; Food R&D Center, Hyundai bioland Co. Ltd. Manhae-ro, Danwon, Ansan, Gyeonggi, Korea

**Keywords:** *Platycodon grandiflorum*, gut microbiota, *Akkermansia*, immune system, diet, microbiome

## Abstract

*Platycodon grandiflorum* (PG) has been used as a traditional remedy to control immune related diseases. However, there is limited information about its immune stimulating effects on the immunosuppressed model. The main bioactive components such as saponins are known to con-tribute to controlling immune activity. Thus, we developed an aged red PG (PGS) with 2.6 times of platycodin D, one of the saponins. We treated PG and PGS (PG-diets) to immunosuppressed mice via cyclophosphamide (CPA) injection. After 2weeks of the supplement, 16S rRNA sequencing was performed to investigate the effects of PG-diets on the gut microbiota and immune system in the immune suppressed model. PG-diets groups showed an increased abundance of microorganism in immune-deficient mice compared to the control NC group, indicating PG-diets have a distinct effect on microbial communities. Detection of specific genera related to the immune related biomarkers in PG-diets groups can support their effects on the immune system. Especially, the *Akkermansia* showed a significant decrease of abundance in response to the CPA treatment in the NC group at the genus level, but its abundance increased in response to the PG-diets treatment in the PG-diets groups. We also found that the modulation of gut microbiome by PG-diets was correlated with body weight as one of important immune biomarkers, though not much difference was found between PG and PGS effects. The results demonstrate that PG-diets may improve the health benefits of immune suppressed mice by altering the gut microbiome.

## INTRODUCTION

Gut microbial community is associated with host digestion, nutrition and even regulation of the host immune system [1]. Microbiota facilitates the development and function of the immune cells at both mucosal and nonmucosal sites. This affects the immune system through the whole body as well as the gut immune system [2]. Given the association between the gut microbiome and immune system, the use of prebiotics is suggested as one of the solutions to improve the immune system. Prebiotics are mostly fibers that are non-digestible food ingredients that can beneficially affect the host’s health by selectively stimulating the activity of some microorganisms [3]. Moreover, prebiotics simulates beneficial bacteria including lactic acid bacteria and bifidobacteria, which increase the expression of anti-inflammatory cytokines and thus enhance the immune system [4].

One of the food ingredients with prebiotic-like effects on the gut microbiota is saponins [5]. Saponin in ginseng (Panax ginseng Meyer) is digested by gut microbiome and the substrate produced by microbiota has effects on the immune system [6]. For instance, ginsenoside, which is a saponin in ginseng, is transformed to compound K by gut microbiota, and the compound K is absorbed into the blood and exhibits potent pharmacological effects such as antitumor, anti-inflammatory, antidiabetic, antiallergic, and neuroprotective effects [7]. The amount of saponin absorption is higher through the metabolism of the gut microbiome than by direct absorption [8]. Also, *Gynostemma pentaphyllum* with abundant saponin boosted the beneficial microbes [9]. Thus, food with enriched saponin is expected to influence the gut microbiome related to the immune system [10].

*Platycodon grandiflorum* (PG), which is known as a herbal medicine in Asia, contains abundant saponin [11]. Specifically, the root of PG has been used to treat various diseases including bronchitis, asthma, pulmonary tuberculosis, diabetes, and inflammatory diseases [12]. The immune-enhancing effect of PG is mainly caused by PG root-derived saponins [13]. In previous studies, PG root-derived saponins were found to have anti-inflammatory and anti-oxidative activity [14]. Furthermore, saponins from PG showed inhibitory effects against infection of Hepatitis C Virus [15]. In our previous in vivo experiment in mice, PG had a role in improving the immune function by increasing body weight, and the serum level of immunoglobulins (IgA and IgM) [16]. IgA and IgM are key factors in the immune system that play a role in the neutralization of toxins, bacteria, or viruses [17]. Recent studies have re-ported that PG increased the level of immunoglobulins in cyclophosphamide (CPA)-immunosuppressed rats, which suggested a positive role in enhancing the immune response [18, 19]. Therefore, the beneficial effects of PG might be linked with an altered gut microbiome, considering saponin is associated with the gut microbiome.

In addition, we developed an aged red PG (PGS) with 2.6 times of platycodin D, which is known as a functional compound in the PG, by steaming and drying the PG. PGS showed improved immune-stimulating effects on the immune-suppressed mice in the previous study [20]. In many studies, functional foods affect the host health by improving the microbiome system. However, there is less study or in-formation about PGS or PG (PG-diets) on microbiome changes in the immune-suppressed mice. Thus, this study was conducted to examine its effects on gut microbiota and immune system in the immune-suppressed mice that are induced with CPA [21]. To validate the effect of PG-diets on the immune system and its association with the gut microbiome, we supplemented PGS and PG to immune-deficient mice. To compare the effect of PG-diets with that of commercial immune-improvement additive, one group was fed by β-glucan [22].

The objective of this study is to reveal the impact of PG-diets with respect to altered microbiota and the immune system. We investigated the changes in the compositions of the gut microbiome in mice exposed to PG-diets and examined the specific genera related to immune-related biomarkers. The outputs can be used to find a functional compound related to enhancing immunity as well as a new functional food material that can contribute to improving the value of domestic PG-diets.

## MATERIALS AND METHODS

### Preparations of test materials

The normal *Platycodon grandiflorum* (PG) and aged PG (PGS) extraction used in this experiment were provided by Hyundai Bioland (Ansan, Gyeonggi, Korea), which is strictly managed and produced according to production management standards. For PGS, it was prepared by washing the roots of domestic PG (Gumsan, Chungnam, Korea) twice, steaming them for 120 minutes, drying them for 24 h, repeating the steaming for 90 min 4 times, and drying them for 72 h to produce the red PGS. The weight of PGS or PG and 50% alcohol were added 15 times compared to the compounds and extracted at 80 ° C for 8 h. The primary extract was recovered, and the remaining underwent secondary extraction with 50% alcohol at 80 ° C for 8 h. All extracts were mixed and filtered using filter press. It was concentrated under reduced pressure at a concentration of 60%, sterilized, and spray dried for the study. PGS and PG extracts were stored at 4°C to protect from light and degradation until use.

### Animals and treatments

Six-week-old C57BL/6 male mice were supplied by Semtaco Co (Chunbuk, Korea). The mice were housed 2 per polycarbonate cage in controlled conditions (20–25°C, 50–55% humidity, and a 12 h light/dark cycle) with free access to water and standard rodent chow (38057, Purinafeed, Gyeonggi-do, Korea). After acclimation for 7 days, a total of 26 mice were randomly divided into seven groups: 1) normal control (Nor); 2) cyclophosphamide (CPA) control (NC); 3) CPA + 2 mg β-glucan/body weight (PC) as a positive control of the preventive treatment; 4) CPA + 75 mg/kg body weight of PGS (PGS1); 5) CPA + 150 mg/kg body weight of PGS (PGS2); 6) CPA + 75 mg/kg body weight of PG (PG1); and 7) CPA + 150 mg/kg body weight of PG (PG2). The experimental extracts dissolved in distilled water were orally administered every day for 2 weeks. Mice of Nor and NC groups were administered an equal volume of distilled water. Immunosuppression was induced by two intraperitoneal injections of CPA (Sigma-Aldrich, St. Louis, MO, USA). CPA was dissolved in saline and 150 mg and 110 mg/kg of CPA were injected intraperitoneally 3 days and 1 day before treatment with PGS or PG, respectively, while the Nor group was injected with an equal volume of saline.

### Metagenome Sequencing

Fecal samples were collected from each mouse and used for DNA extraction using AccuPrep Stool DNA Extraction Kit following the instructions of the manufacturer. After performing quality control (QC), qualified samples were proceeded to library construction. The V3 and V4 region of the 16S rRNA genes were PCR amplified from the microbial ge-nomic DNA. The DNA quality was determined by PicoGreen and Nanodrop. The input gDNA (10 ng) was PCR amplified using the barcoded fusion primers 341F/805R (341F: 5 ′ CCTACGGGNGGCWGCAG 3, 805R: 5 ′ GACTACHVGGGTATCTAATCC 3′). The final purified product was quantified using qPCR according to the qPCR Quantification Proto-col Guide (KAPA Library Quantification kits for Illumina Sequencing platforms) and qualified using the LabChip GX HT DNA High Sensitivity Kit (PerkinElmer, Massachusetts, USA). The 300 paired-end sequencing reaction was performed on the MiSeq™ platform (Illumina, San Diego, USA). The sequencing data generated for this study are available at the Sequence Read Archive (SRA) of NCBI (http://www.ncbi.nlm.nih.gov/sra) under BioProject PRJNA700675.

### Raw Data Processing and Taxonomic Analysis

Pre-processed reads were imported and analyzed using QIIME2 version 2020.02 [23]. We used the DADA2 software package implemented in QIIME2 to denoise and correct Illumina sequenced FASTA2Q files by removing chimeras sequences using the “consensus” method [24]. Based on the quality plot of the DADA2, reads were trimmed using the following parameters: -p-trunc-len-f=300; -p-trunc-len-r=240; -p-trim-left-f=6; and -p-trim-left-r=6. For taxonomy assignment, a naïve Bayes classifier was trained on a GreenGenes 97% (version 13.8) operational taxonomic unit (OTU) database with reference sequences trimmed to the V3-V4 region using QIIME2 plugin feature-classifier [25].

### Diversity Analysis

All the sequence data were rarefied to a sampling depth of 12,361 sequences per sample prior to computation of alpha and beta-diversity analysis with QIIME2 plugin diversity, such as observed OTUs, Shannon, and weighted Unifrac. The weighted Unifrac distance matrix was used for nonparametric Permutation Multivariate Analysis of Variance (PERMANOVA) and Principal Coordinate Analysis (PCoA) plot [26-29]. PER-MANOVA was performed with 999 permutations to weighted UniFrac distance matrix using Adonis function in R package ‘vegan’ [30].

### Performance Analysis

In the present study, body weight and Serum immunoglobulin measurements of e-perimental mice were measured. A total of 26 samples (three or four mice in each of the seven groups) similar to the average group weight were selected for the performance analysis. Body weights were monitored once a week. All animals were overnight fasted (water was not restricted) before initial test substance administration and sacrifice to re-duce the individual differences from feeding. The mice were sacrificed under inhalation anesthetized with CO2, using rodent inhalation anesthesia apparatus (Surgivet, Waukesha, WI, USA). Serum concentrations of immunoglobulin A (IgA) and immuno-globulin M (IgM) were measured using enzyme-linked immunosorbent assays kits (ELISA; Abcam, USA), according to the manufacturer’s instructions. All the assays were performed in duplicate. All animals were treated according to the international regulations for the usage and welfare of laboratory animals. This study was approved by the Institutional Animal Care and Use Committee in the National Institute of Agricultural Sciences (NAS-201808).

### Statistical analysis

Trimmed Mean of M values (TMM) was obtained to adjust for different library sizes using edgeR [31]. Then, statistical tests were performed under a generalized linear model (GLM) considering OTU’s count as a negative binomial distribution. To compare the good-ness-of-fit of two models, the log-likelihood ratio statistic was calculated, and the false discovery rate (FDR) was used to adjust for multiple testing errors with a significance level of 5% [32]. Another statistical test for differentially abundant microbial taxa was assessed using the QIIME2 Analysis of Composition of Microbiomes (ANCOM) plugin in order to identify features that significantly differed in abundance from each study group [33]. The differentially abundant features at each phylogenetic level were calculated by ANCOM from the DADA2 feature table. The Student’s t-test was used to compare the biomarkers (growth weight, IgA, and IgM) between groups. Also, the pair-wise correlations between the abundance of microbiota and immune-related biomarkers were determined using Spear-man correlation coefficient (r) from the *corplot* R package. The abundance of significantly correlated genera (p-value < 0.05) was visualized in a line graph with the corresponding phenotype to see its correlations. All R packages were implemented in RStudio version 4.0.1 [34].

### Functional Profiles of Gut Microbiota

Phylogenetic Investigation of Communities by Reconstruction of Unobserved States2 (PICRUSt2) was used to predict Kyoto Encyclopedia of Genes and Genomes (KEGG) Orthology (KO) genes from 16S rRNA data [35]. The tree generated in PICRUSt2 with the maximum nearest-sequenced taxon index (NSTI) cut-off of 2 was complemented with the feature table resulted from QIIME2 plugin dada2 for hidden-state prediction (hsp) using the maximum-parsimony method [36]. Followed by the prediction of KO genes, the KEGG pathways were then mapped to the KO genes using the PICRUSt2 package. Afterward, multiple group comparison was computed using the Analysis of Variance (ANOVA) statistical test (p-value<0.05) in the STAMP software package to carry out significant pathways [37]. Moreover, functions related to human diseases were achieved using White’s non-parametric t-test from the STAMP software package [38].

## RESULTS

### Effects of *Platycodon grandiflorum* on host immune system and gut microbiome diversity

The principal coordinate analysis (PCoA) with weighted Unifrac distance was per-formed in two comparisons: 1) between normal control (Nor) group and cyclophosphamide (CPA) control as a negative control (NC) group, and 2) between NC and groups supplemented by β-glucan (PC), PGS, and PG. In the first comparison, Nor and NC groups were segregated but they had high variance within groups (Figure 1A). When we compared NC to other groups that are immunosuppressed with CPA, the positive control with β-glucan (PC) and PG2 groups had no segregation with the NC group (Figure 1B). On the other hand, PGS1, PGS2, and PG1 groups are distinguished from the NC group, where the PGS2 group was clustered at a shorter distance compared with those in the other groups.

**Figure 1.**
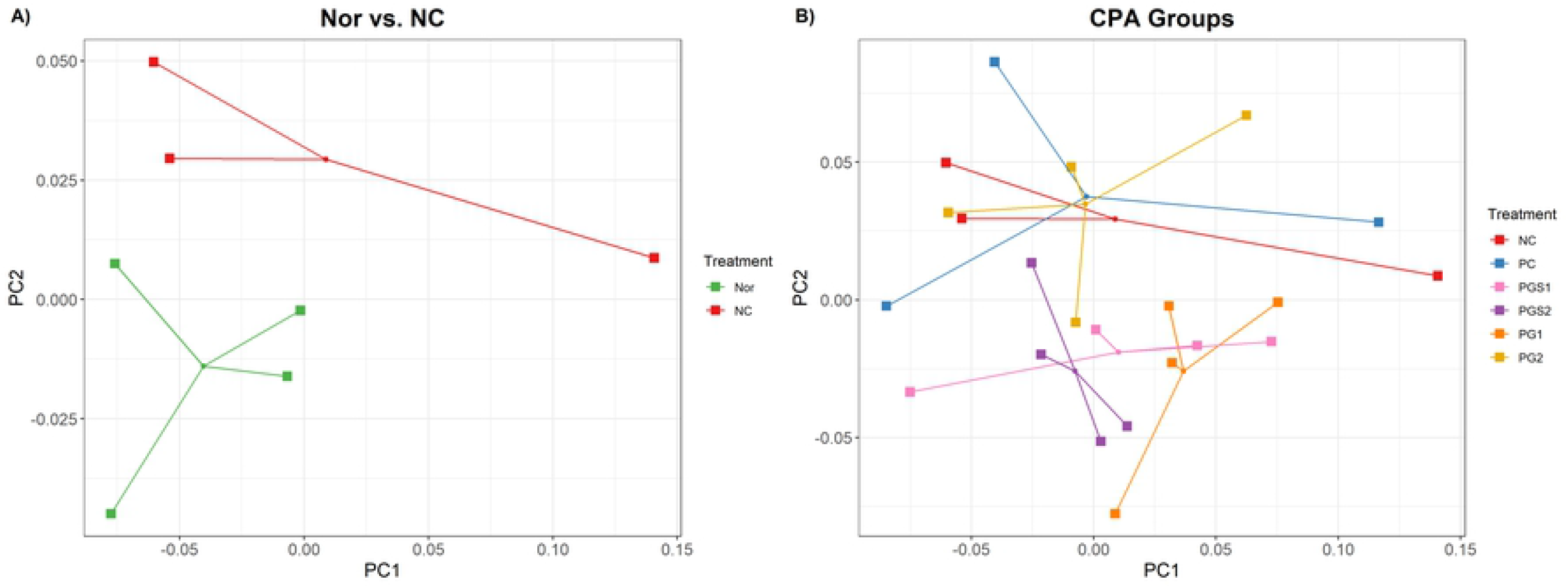
A principal coordinate analysis (PCoA) plot showing dissimilarities among different diet groups. **A**) Comparison between Nor and NC using PCoA from the distance of weighted UniFrac. Each dot represents one sample. Green dots are normal control group (Nor) and red dots are negative control group (NC) that are immunosuppressed with cyclophosphamide (CPA). **B**) Comparison of immunosuppressed with CPA groups named NC, positive control (PC), aged red PG (PGS1, PGS2), and PG (PG1, PG2) using PCoA plot.

Along with PCoA, a nonparametric Permutation Multivariate Analysis of Variance (PERMANOVA) was performed with 999 permutations and used to determine the significant differences between the seven groups. Pairwise combinations with PG1 have shown significantly lower p-values than that of the other pairwise combinations, indicating that the abundance and variety of microorganisms in immunodeficient mice exposed to PG1 differed from that of the NC and PC groups. This implies that this can alter the composition similar to the healthy Nor group.

In terms of alpha-diversity, microbial diversity within a local community was evaluated based on species richness and diversity using the observed operational taxonomic units (OTUs) and Shannon index. Among the seven groups, there was no significant difference found in both richness and diversity, but they showed a similar distribution across combinations (S1 Fig).

### Change of microorganism abundance by supplement of *Platycodon grandiflorum*

The predominant microbes at the phylum level were Bacteroidetes and Firmicutes. They account for more than 70% and 20% of microbes (Fig 2A), respectively. The Bacteroidetes, when observed in the NC group, showed a decrease in abundance but an increase in PG-diets. Alongside, the *Firmicutes* showed a substantial increase in abundance observed in PG-diets. Since only limited information was available regarding the differences at the phylum level, we compared the abundance of the NC group to those of PC, PGS, and PG treated groups to detect supplement of PGS associated microorganisms at the genus level. By using the weighted trimmed mean of the log expression ratios (trimmed mean of M-values (TMM)), NC group associated microorganisms were *Akkermansia* and *Staphylococcus* (FDR < 0.05) (Fig 2B). The abundance of *Akkermansia* decreased in the NC group compared to the Nor group. When immune suppressed mice had a supplement with PG-diets, the abundance of *Akkermansia* increased. There were significant differences within the NC group and PGS2 and PG1 groups (p-value < 0.05). On the other hand, the abundance of Staphylococcus increased in the NC group compared to the Nor group. Unlike *Akkermansia*, there was no distinct difference within the groups, but decrease in *Staphylococcus* abundance was observed in PC and PG-diets groups. We also carried out the Analysis of Composition of Microbiomes (ANCOM) implemented through the QIIME2 ANCOM plugin to investigate differentially abundant microorganisms at phylum and order levels and compare with the results of edgeR. There were nine phyla identified but *Verrucomicrobia* were the only phylum to demonstrate a significant change in response to treatments. Also, this observation was consistent for the taxonomic classification levels of order, such that the order of *Verrucomicrobiales* was significantly different among treatments (S2 Fig).

**Figure 2.**
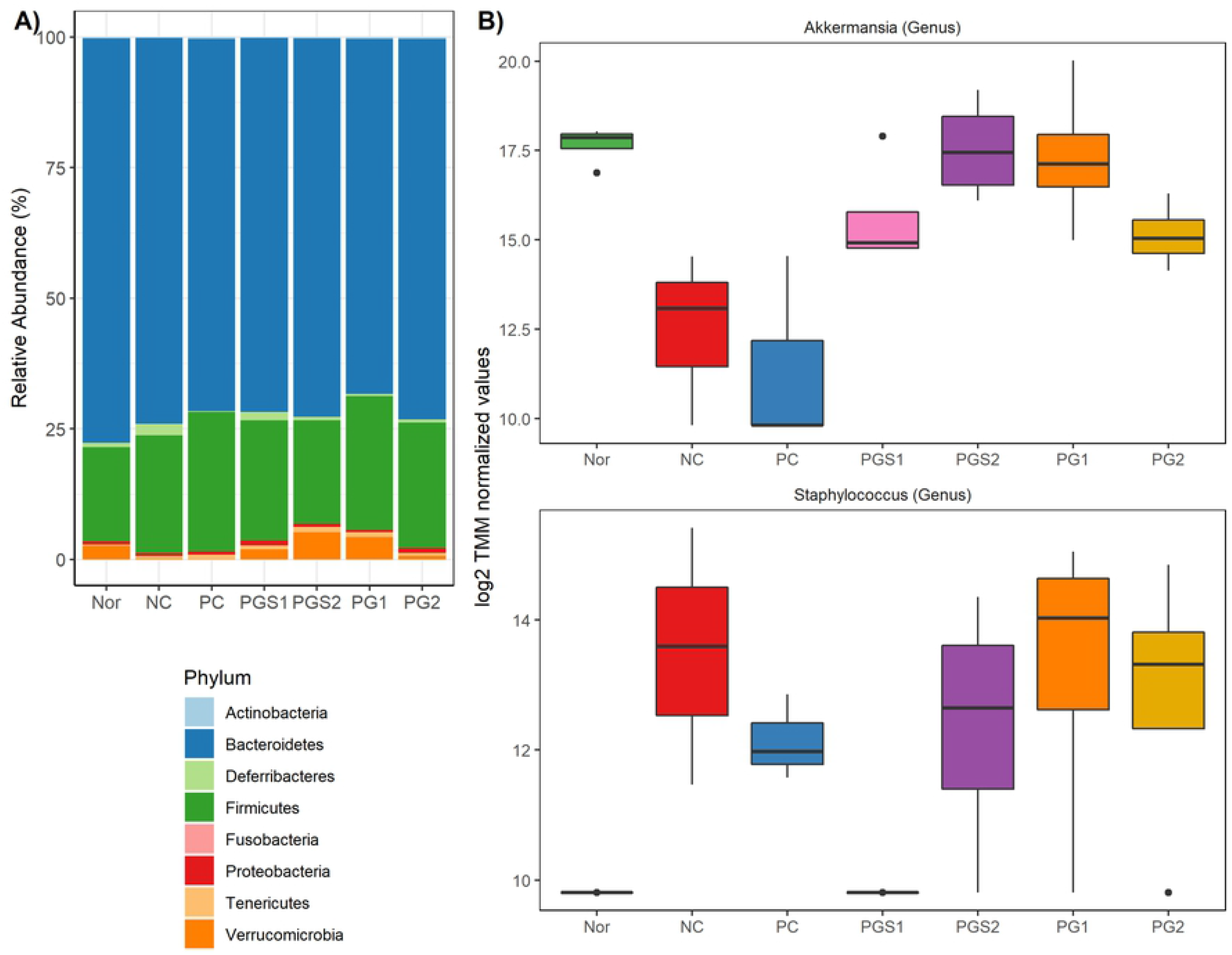
Relative Abundance of microorganisms at phylum and genus levels. **A**) Phylum level composition. Bar plots represent the percentage (%) of average abundance for each group. **B**) Differentially abundant values among genus level between control and immune suppress groups. Two genera (*Akkermansia* and *Staphylococcus*) were differentially abundant with a significant FDR<0.05.

### General characteristics of phenotypes in seven groups

Body weights and serum immunoglobulin (IgA and IgM) levels for seven groups were measured in the present study. The Nor group had the highest body weight and IgA (Fig 3A,B). For body weight, NC group showed a significant decrease compared to the Nor group (p-value=0.0103). PG1 group showed a significant increase compared to the NC group (p-value=0.049). Body weight increased in PGS1 and PGS1 groups compared to the NC group (p>0.05). A similar observation was found in IgA, where the NC group showed a significantly lower IgA value than the Nor group (p-value=0.0156). The IgA levels of PGS1, PGS2, and PG2 were higher than that of the NC group (p>0.05). Likewise, PG-diets groups showed higher serum IgM levels compared to the NC group (p>0.05; Fig 3C). Thus, PG-diets significantly affect body weight in the immune-suppressed mice and tend to increase levels of IgA and IgM, but not much of clear significant differences were found between NC and PG-diets or within PG-diets groups.

**Figure 3.**
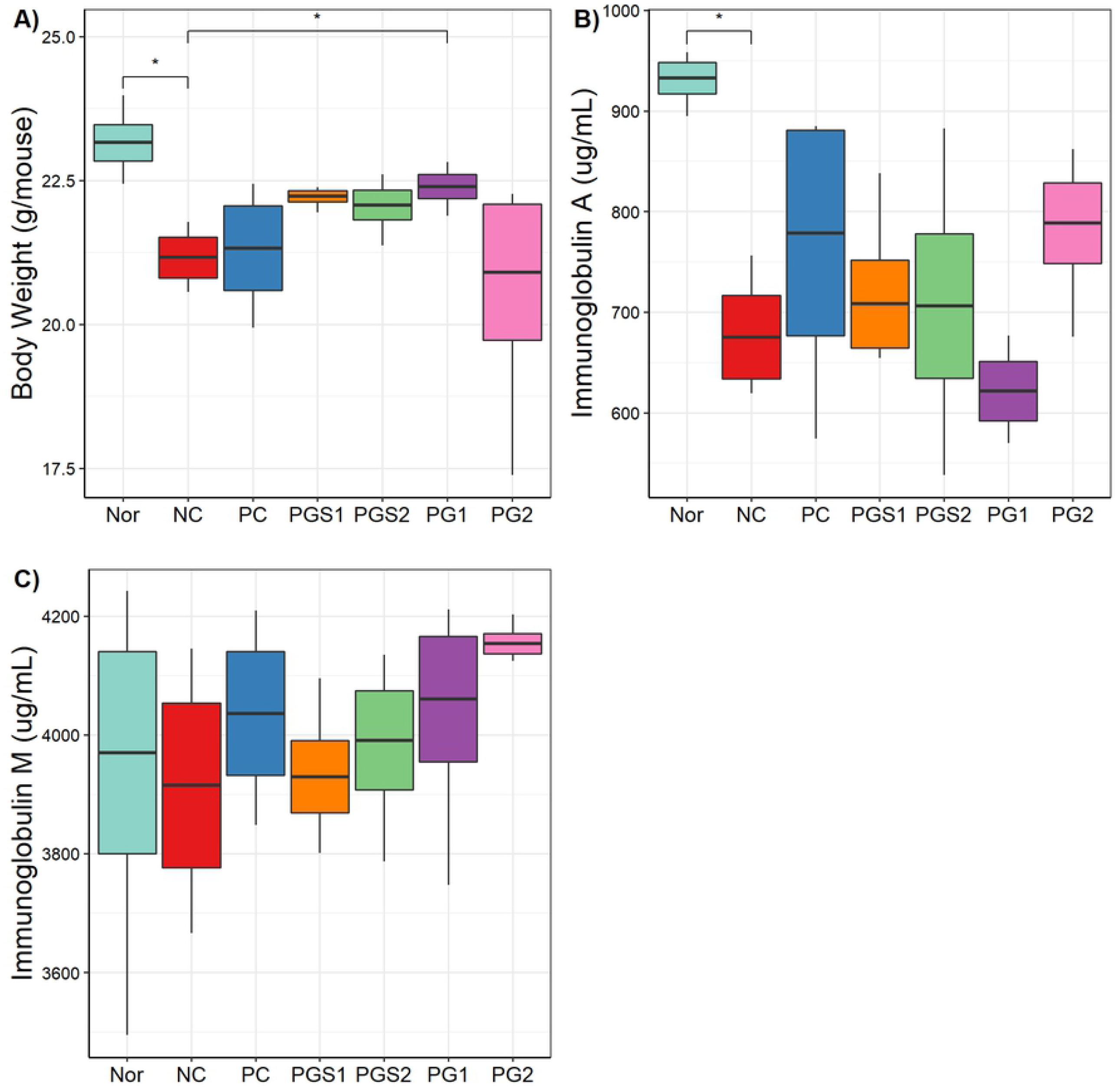
Body weight and serum levels of immunoglobulins (IgA, IgM) of mice. Nor group showed a higher body weight and IgA. Immune suppressed mice (NC) decreased in body weight, IgA and IgM compared to Nor group and increasing observations were shown in supplementary effects of PG-diets. (*: significance with p-value<0.05 cutoff).

### Correlation between the immune related biomarkers and relative abundance of microorganisms

P. grandiflorum has shown to affect the body weight, one of the immune-related biomarkers, and the microbiome composition such as *Akkermansia* genus, *Lactobacillaceae* family, *Gemellates* order, and *Staphylococcus* genus (Fig 4). Spearman’s rank correlation coefficient was significant for four microorganisms (Spearman’s rho; p-value<0.05). Among the four genera associated with body weight, *Akkermansia* showed the highest positive correlation, and *Staphylococcus* showed a highly negative correlation with body weight, given the 0.54 and −0.43 of Spearman’s rho, respectively (Fig 4). Moreover, six microorganisms were associated with IgA level, with three species found to be positively correlated and three species were negatively correlated (S3 Fig A). In IgM level, three species were positively correlated, and rest was negatively correlated (S3 Fig B). If a positive correlation is present, the abundance of micro-organism tends to increase as the body weight increases. Vice versa with the negative correlation, the abundance of microorganism decreases as the body weight increases.

**Figure 4.**
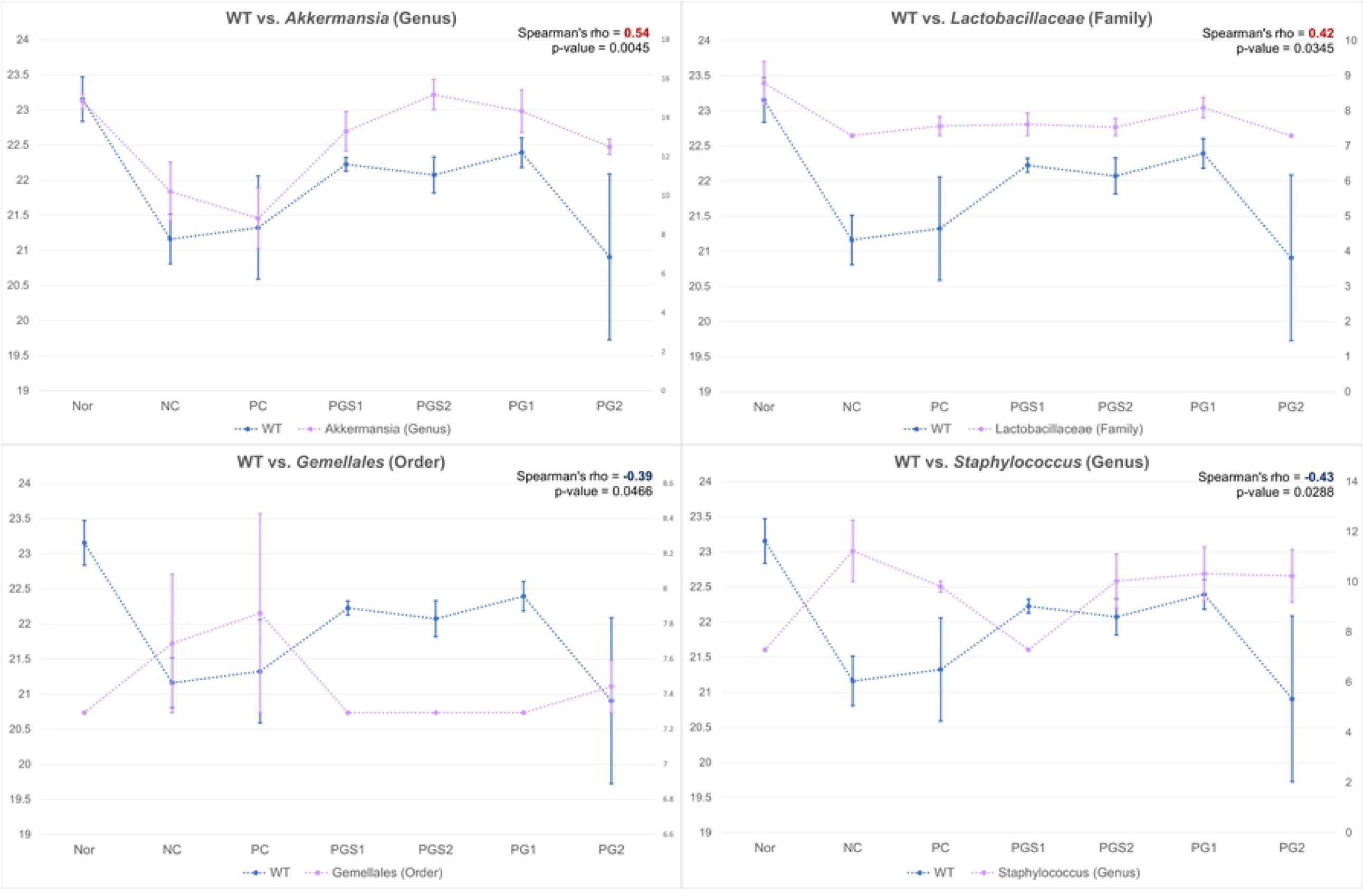
Correlation between immune related biomarkers and relative abundance of microorganisms. Four significant genera were correlated with body weight. The x-axis is the group, and the left y-axis is the body weight value (blue) and right y-axis is the relative abundance value (pink). The Spearman’s rho and the p-value for the correlations were shown on right top corner.

### Functional pathways of the microbiota

We performed Phylogenetic Investigation of Communities by Reconstruction of Unobserved States (PICRUSt2) to predict functional pathway abundance related to PG-diets. Then, multiple groups comparison was computed using the STAMP software and the ANOVA statistical test. Out of 137 Kyoto Encyclopedia of Genes and Genomes (KEGG) pathways, 2 functional pathways were significant in the study (p-value<0.05; Fig 5). NC group showed enriched carbon fixation, whereas PG-diets showed enriched in selenocompound metabolism.

**Figure 5.**
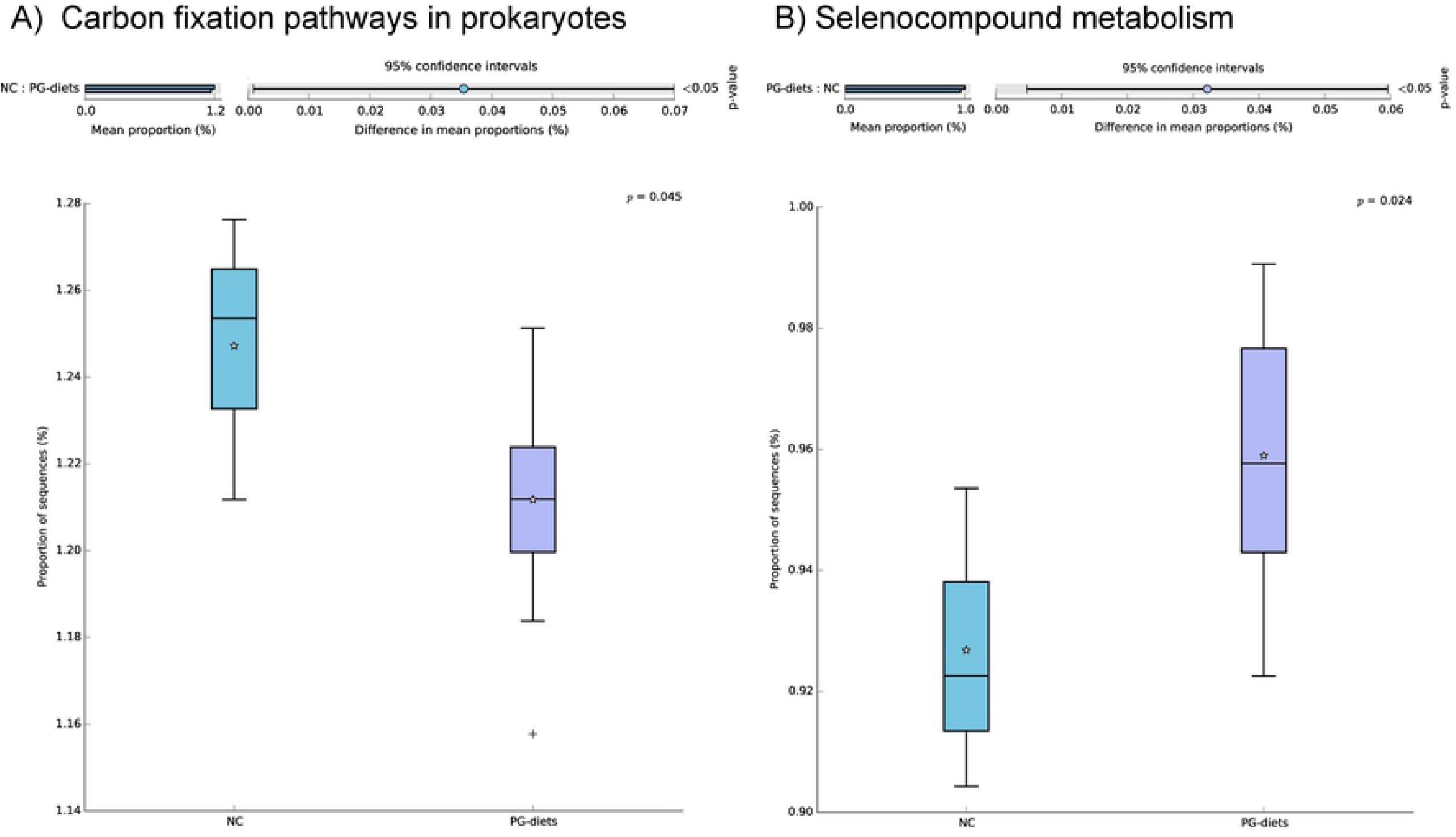
Predicted microbial functions showing significant differences between NC and PG-diet groups. Functions were predicted by PICRUSt2 against the KEGG pathway database and ANOVA statistical test (*p*-value<0.05) was performed using the STAMP software program. **A**) Carbon fixation pathways in prokaryotes were enriched in the NC group. **B**) Selenocompound metabolism was enriched in the PG-diet group.

## DISCUSSION

In this study, we demonstrated the effect of *P. grandiflorum* on the gut microbiome, especially in the immune suppressed mouse models by CPA. The impact of the PG-diets on the altered gut microbiome in this study may support our previous study, which high-lighted the preventive effect of PGS and PG extracts (PG-diets) on the immunosuppressed system in the NC group. As a result of multiple analyses of gut microbiome and phenotype of immune-suppressed mice, our study provides a deeper insight into the relationship between the intake of P. grandiflorum root, microbial communities, and immune-related biomarkers.

The chemical named CPA that we used to suppress the immune system in mice is commonly used for treating repulsions and malignant tumors during organ or bone mar-row transplants. However, CPA appeared to be toxic to normal cells, causing not only side effects such as weight loss, acute leukemia, liver dysfunction, anemia, and hair loss, but also the seriousness of the problems that arise in long-term use [39-41]. Thus, it is imperative to develop natural materials that can suppress the toxic or side effects of toxic immunosuppressants. As the root of *P. grandiflorum* is known to contain a lot of fibers, calcium, iron, and functional compound such as saponin, it is used to treat patients with bronchial diseases such as bronchitis and asthma [42, 43]. In addition, previous *in vivo* experiments reported that the root of PG improved the immunity of immunosuppressed mice at 150mg/kg body weight [16]. However, the PG has a strong bitter taste which urged us develop the aging red PG (PGS) that not only improved the taste, but also the content of a physiologically-active substance called platycodin D. This has anti-cancer effects in addition to anti-inflammatory and diabetes prevention [44]. Moreover, it is well known to improve immunity by proliferating immune cells [45]. Thus, considering the PG with 150mg/kg body weight showed an enhancing immunity in *in vivo* experiment, PGS with 150mg/kg and half of its amount (75mg/kg) were evaluated to characterize the effect of PG. In our study, we revealed that supplement with PG-diets affected the reduced body weights of the mouse due to the CPA treatment, which showed that their body weights recovered to that of the normal group. It also increased the serum level of immunoglobulins (IgA and IgM) in the groups treated with each material without adverse or toxic effects after oral administration.

A decrease in body weight is considered a side effect of long-term use of CPA treatment. As the NC group showed a significant decrease in body weight compared to the Nor group, it then reinforces the side effects of CPA treatment. From the statistical t-test, the PG1 group showed a significantly increased body weight. Although PGS1 and PGS2 did not show a significant increase compared to the NC group, they displayed higher body weight than those of the NC and PC groups.

In our *in vivo* study, PG-diets increased body weight decreased due to CPA, and a significant effect was found in the PG1 group compared to the NC group, which agrees with the observations in previous reports [16, 20]. In addition, we observed that the concentration of IgA significantly reduced in the NC group compared to the Nor group. Though, their levels of PGS1, PGS2 and PG2 groups increased compared to the Nor group (p>0.05). A similar observation was shown in IgM where the PG-diets groups leveled higher than the NC group. Therefore, our data suggest that CPA caused a significant reduction in body weight and immunoglobulin levels by its toxicity while PG-diets can prevent the side effects of CPA and can improve the conditions of mice treated with PG-diets. These findings indicate that the PG-diets can resist the immunosuppressive effects of CPA.

The gut microbiome diversity of seven groups has been shown to elucidate the significant differences between groups. The microbiota in the Nor and NC groups have differed and the PG-diets groups, except for the PG2 group, were segregated with respect to the NC group, thereby indicating that there is a distinct diversity between them. The altered abundance of bacteria following the PG-diets occurs at the generic level. The *Akkermansia* appeared to increase in the PG-diets. Just like the observation shown in body weight, the NC group decreased the abundance of *Akkermansia* compared to the Nor group. Although the PC group which supplemented with β-glucan shows a weak effect on the abundance, PG-diets increased to a level similar to the Nor group. β-glucan is known to play a role in enhancing the immune system without becoming overactive, lowering elevated levels of low-density lipoprotein (LDL) cholesterol, and helping prevent infections as well as the prevention and treatment of cancer [46, 47]. Interestingly, PG-diets may be more effective than β-glucan though small sample sizes (n=3∼4) used for this experiment have some limitation. Moreover, previous papers reported that an increase in *Akkermansia* leads to anti-inflammatory activity in the intestinal tract and reduces diabetes [48, 49]. It is also introduced as beneficial bacteria. These have supported our results where we presented a positive correlation with body weight and *Akkermansia*. Another significant microbe shown at the generic level was *Staphylococcus*. NC group increased the level compared to the Nor group while the PGS groups decreased it though a clear difference was not shown between PG-diets. *Staphylococcus* is known to cause food poisoning or skin allergy by its toxin [2]. Thus, the lower the abundance indicates the healthier the gut. It also displayed a negative correlation with body weight. Although the underlying mechanisms are unknown, these results support the idea of PG-diets altering the gut microbiome.

The association between microbiome and other traits including IgA and IgM levels was also determined, which represents candidate microorganisms of its traits. Immuno-globulin levels were investigated as an index of the immune system related to the microbiome [50]. The immunoglobulin levels of PG-diets in *in vivo* experiment were generally higher than that of the NC group (p>0.05). Moreover, the genera associated with immunoglobulin levels were consistent with or contrary to previous metagenome studies related to the immune system. *Ruminococcaceae* and *Clostridium* groups that were negatively and positively correlated with IgA, respectively, were also involved in inflammatory diseases. As *Ruminococcaceae* worked in the inoculation of strongly virulent inflammatory bowel disease, the result of lower IgA level could be related to more microorganism. In addition, functional analysis revealed that a combination of PG-diets was enriched in selenocompound metabolism. Selenium (Se) is essential micronutrient for animals’ metabolism and is required for the biosynthesis of selenoproteins, which participate in the immune response, cancer chemoprevention, and other processes [51]. Therefore, the correlation between IgA level and enrichment of selenocompound metabolism suggests that altered communities of microorganisms induced by PG may alter the levels of immunoglobulin, and can be linked to the regulation of the immune system.

Our results showed the degree of alternation based on the supplementation of PG in immunodeficiency. Although the small sample size and amount of intake were the study limitations, this was the first study to investigate the effect of PG on the gut microbiome and immune system, despite several studies reporting its health benefits. The abundance of *Akkermansia* was increased in immunodeficient mouse supplemented with PG-diets, indicating that the PG has a distinct effect on microbial communities. Furthermore, we investigated specific genera related to body weight and serum immunoglobulin levels, which were known to be important for the immune system. As PG-diets have provided a functional compound related to enhancing immunity, the health benefit of PG-diets in immunodeficiency may mediate via its microbiome, such as *Akkermansia* microorganism. Moreover, PG-diets could be a potential nourishing immunity supplement for an immunosuppressed model. In addition, PG-diets could contribute to improving the value of its domestic PG, as well as enhancing the reliability of domestic foods.

## Supporting Information

**S1 Fig. Observed OTUs and Shannon index in seven groups**. In observed OTUs, richness tends to increase in PG-diets groups compared to NC group. However, no clear significant differences were shown in both richness and diversity.

**S2 Fig. Differentially abundant microbial taxa identified by q2-ANCOM**. ANCOM volcano plot of differential abundance at the group level, where the x-axis represents F-statistics and y-axis represents the W-statistics. The F-statistics are a measure of the effect size difference for a particular species between the study groups and the W-statistic is the strength of the ANCOM test for the tested number of species. A) At the phylum level, the square represents *Verrucomicrobia*, the triangle represents *Bacteria, Actinobacteria, Bacteroidetes, Firmicutes, Fusobacteria, Proteobacteria, and Tenericutes*, circle represents *Deferribacteres*. B) At the order level, the square represents *Verrucomicrobiales*, the triangle represents *Bacteroidales, Bacillales, Gemellales, Lactobacillales, Clostridiales, Fusobacteriales, Pseudomonadales*, and *RF39*, circle represents *Actinomycetales, Coriobacteriales, Deferribacterales, Turicibacterales, Erysipelotrichales, Burkholderiales, Desulfovibrionales*, and *Anaeroplasmatales*.

**S3 Fig. Correlation between IgA and IgM level and relative abundance of microorganisms**. Total of 6 and 5 significant genera were correlated with IgA and IgM, respectively (A, B). The x-axis is the group, and the left y-axis is the phenotype values (blue) and right y-axis is the relative abundance values (IgA: yellow; IgM: orange). The Spearman’s coefficient and p-value are placed on the right top corner.

## Acknowledgements

The authors appreciate Mara Shyn Valdeabella for reviewing the manuscript, and Jung Hyun Lim and Min Sook Kim for their supporting the animal experiment. We also thanks eGnome for helping sample analysis.

## Author Contributions

Conceptualization: Sung-Hyen Lee

Methodology: Sung-Hyen Lee, So Yun Jhang, Hyun Cheol Jeong, Sung Jin Lee, Hyun-Ju Kim

Data Validation: Sung-Hyen Lee, So Yun Jhang, Sohyun Bang

Formal analysis: Sung-Hyen Lee, So Yun Jhang, Sohyun Bang

Investigation: Sung-Hyen Lee

Resources: Sung-Hyen Lee, Eun-Byeol Lee, Ji-Hye Choi, Hae-Jeung Lee

Writing – original draft: Sung-Hyen Lee, So Yun Jhang

Writing, review & editing: Sung-Hyen Lee, So Yun Jhang, Sohyun Bang, Hwan-Hee Jang

Supervision, project administration and funding acquisition: Sung-Hyen Lee

## Notes

### Competing Interest Statement

The authors have declared no competing interest.

## Reference

1. RooksBelkaid Y, Hand TW. Role of the microbiota in immunity and inflammation. Cell. 2014;157(1):121–41.

2. Belkaid Y, Hand TW. Role of the microbiota in immunity and inflammation. Cell. 2014;157(1):12-41.

3. Schley P, Field C. The immune-enhancing effects of dietary fibres and prebiotics. British Journal of Nutrition. 2002;87(S2):S221–S30.

4. Lomax AR, Calder PC. Prebiotics, immune function, infection and inflammation: a review of the evidence. British Journal of nutrition. 2008;101(5):633–58.

5. Chen L, Tai WC, Hsiao WW. Dietary saponins from four popular herbal tea exert prebiotic-like effects on gut microbiota in C57BL/6 mice. Journal of Functional Foods. 2015;17:892–902.

6. Lee J, Lee E, Kim D, Lee J, Yoo J, Koh B. Studies on absorption, distribution and metabolism of ginseng in humans after oral administration. Journal of ethnopharmacology. 2009;122(1):143–8.

7. Li Y, Zhou T, Ma C, Song W, Zhang J, Yu Z. Ginsenoside metabolite compound K enhances the efficacy of cisplatin in lung cancer cells. Journal of thoracic disease. 2015;7(3):400.

8. Kim K-A, Yoo HH, Gu W, Yu D-H, Jin MJ, Choi H-L, et al. A prebiotic fiber increases the formation and subsequent absorption of compound K following oral administration of ginseng in rats. Journal of ginseng research. 2015;39(2):183–7.

9. Chen L, Brar MS, Leung FC, Hsiao WW. Triterpenoid herbal saponins enhance beneficial bacteria, decrease sulfate-reducing bacteria, modulate inflammatory intestinal microenvironment and exert cancer preventive effects in ApcMin/+ mice. Oncotarget. 2016;7(21):31226.

10. Kim D-H. Gut microbiota-mediated pharmacokinetics of ginseng saponins. Journal of ginseng research. 2018;42(3):255–63.

11. Han L-K, Xu B-J, Kimura Y, Zheng Y-n, Okuda H. Platycodi radix affects lipid metabolism in mice with high fat diet–induced obesity. The Journal of nutrition. 2000;130(11):2760–4.

12. Lee EB. Pharmacological studies on Platycodon grandiflorum A. DC. IV. A comparison of experimental pharmacological effects of crude playtcodin with clinical indications of platycodi radix (author’s transl). Journal of the Pharmaceutical Society of Japan. 1973;93(9):1188–94.

13. Lee BJ, Jeon SH, No IR, Kim YG, Cho YS. Effect of saponin content and antioxidant activities of Platycodon grandiflorum Radix by cutting length. Korean Journal of Medicinal Crop Science. 2015;23(5):363–9.

14. Lee KJ, Choi CY, Chung YC, Kim YS, Ryu SY, Roh SH, et al. Protective effect of saponins derived from roots of Platycodon grandiflorum on tert-butyl hydroperoxide-induced oxidative hepatotoxicity. Toxicology letters. 2004;147(3):271–82.

15. Kim J-W, Park SJ, Lim JH, Yang JW, Shin JC, Lee SW, et al. Triterpenoid saponins isolated from Platycodon grandiflorum inhibit hepatitis C virus replication. Evidence-Based Complementary and Alternative Medicine. 2013;2013.

16. Lee EB, Lee SH, Park Y-G, Choi J-H, Lee HK, Jang HH, et al. Platycodon grandiflorum Extract Ameliorates Cyclophosphamide-Induced Immunosuppression in Mice. J East Asian Soc Diet Life. 2019;29(4):303-9. doi: https://doi.org/10.17495/easdl.2019.8.29.4.303.

17. Schroeder HW, Jr., Cavacini L. Structure and function of immunoglobulins. J Allergy Clin Immunol. 2010;125(2 Suppl 2):S41–52. Epub 2010/03/05. doi: 10.1016/j.jaci.2009.09.046. PubMed PMID: 20176268; PubMed Central PMCID: PMCPMC3670108.

18. Noh EM, Kim JM, Lee HY, Song HK, Joung SO, Yang HJ, et al. Immuno-enhancement effects of Platycodon grandiflorum extracts in splenocytes and a cyclophosphamide-induced immunosuppressed rat model. BMC Complement Altern Med. 2019;19(1):322. Epub 2019/11/23. doi: 10.1186/s12906-019-2724-0. PubMed PMID: 31752816; PubMed Central PMCID: PMCPMC6868875.

19. Yu Q, Nie SP, Wang JQ, Liu XZ, Yin PF, Huang DF, et al. Chemoprotective effects of Ganoderma atrum polysaccharide in cyclophosphamide-induced mice. Int J Biol Macromol. 2014;64:395–401. Epub 2013/12/29. doi: 10.1016/j.ijbiomac.2013.12.029. PubMed PMID: 24370474.

20. Choi J-H, Lee EB, Park Y-G, Lee HK, Jang HH, Choe J, et al. Aged Doraji (Platycodon grandiflorum) Ameliorates Cyclophosphamide-Induced Immunosuppression in Mice. The Korean Society of Pharmacognosy. 2019;50(3):219–25.

21. Zhou Y, Chen X, Yi R, Li G, Sun P, Qian Y, et al. Immunomodulatory effect of tremella polysaccharides against cyclophosphamide-induced immunosuppression in mice. Molecules. 2018;23(2):239.

22. Sandvik A, Wang Y, Morton H, Aasen A, Wang J, Johansen FE. Oral and systemic administration of β-glucan protects against lipopolysaccharide-induced shock and organ injury in rats. Clinical & Experimental Immunology. 2007;148(1):168–77.

23. Bolyen E, Rideout JR, Dillon MR, Bokulich NA, Abnet CC, Al-Ghalith GA, et al. Reproducible, interactive, scalable and extensible microbiome data science using QIIME 2. Nat Biotechnol. 2019;37(8):852–7. Epub 2019/07/26. doi: 10.1038/s41587-019-0209-9. PubMed PMID: 31341288; PubMed Central PMCID: PMCPMC7015180.

24. Callahan BJ, McMurdie PJ, Rosen MJ, Han AW, Johnson AJ, Holmes SP. DADA2: High-resolution sample inference from Illumina amplicon data. Nature methods. 2016;13(7):581–3. doi: 10.1038/nmeth.3869. PubMed PMID: 27214047; PubMed Central PMCID: PMC4927377.

25. McDonald D, Price MN, Goodrich J, Nawrocki EP, DeSantis TZ, Probst A, et al. An improved Greengenes taxonomy with explicit ranks for ecological and evolutionary analyses of bacteria and archaea. ISME J. 2012;6(3):610–8. Epub 2011/12/03. doi: 10.1038/ismej.2011.139. PubMed PMID: 22134646; PubMed Central PMCID: PMCPMC3280142.

26. Lozupone C, Knight R. UniFrac: a new phylogenetic method for comparing microbial communities. Appl Environ Microbiol. 2005;71(12):8228–35. Epub 2005/12/08. doi: 10.1128/AEM.71.12.8228-8235.2005. PubMed PMID: 16332807; PubMed Central PMCID: PMCPMC1317376.

27. Lozupone C, Lladser ME, Knights D, Stombaugh J, Knight R. UniFrac: an effective distance metric for microbial community comparison. ISME J. 2011;5(2):169–72. Epub 2010/09/10. doi: 10.1038/ismej.2010.133. PubMed PMID: 20827291; PubMed Central PMCID: PMCPMC3105689.

28. Chang Q, Luan Y, Sun F. Variance adjusted weighted UniFrac: a powerful beta diversity measure for comparing communities based on phylogeny. BMC Bioinformatics. 2011;12:118. Epub 2011/04/27. doi: 10.1186/1471-2105-12-118. PubMed PMID: 21518444; PubMed Central PMCID: PMCPMC3108311.

29. Noma H, Nagashima K, Furukawa TA. Permutation inference methods for multivariate meta-analysis. Biometrics. 2020;76(1):337–47. Epub 2019/08/11. doi: 10.1111/biom.13134. PubMed PMID: 31399994.

30. Oksanen JB, F. Guillaume; Friendly, Michael; Kindt, Roeland; Legendre, Pierre; McGlinn, Dan; Minchin, R. Peter; O’Hara, R.B.; Simpson, Gavin L.; Solymos, Peter; Stevens, M. Henry H.; Szoeces, Eduard; Wagner, Helene. vegan: Community Ecology Package. 2019.

31. Robinson MD, McCarthy DJ, Smyth GK. edgeR: a Bioconductor package for differential expression analysis of digital gene expression data. Bioinformatics. 2010;26(1):139–40.

32. Benjamini Y, Hochberg Y. Controlling the false discovery rate: a practical and powerful approach to multiple testing. Journal of the royal statistical society Series B (Methodological). 1995:289–300.

33. Mandal S, Van Treuren W, White RA, Eggesbo M, Knight R, Peddada SD. Analysis of composition of microbiomes: a novel method for studying microbial composition. Microb Ecol Health Dis. 2015;26:27663. Epub 2015/06/02. doi: 10.3402/mehd.v26.27663. PubMed PMID: 26028277; PubMed Central PMCID: PMCPMC4450248.

34. Team R. RStudio: Integrated Development Environment for R. 2020.

35. Langille MG, Zaneveld J, Caporaso JG, McDonald D, Knights D, Reyes JA, et al. Predictive functional profiling of microbial communities using 16S rRNA marker gene sequences. Nat Biotechnol. 2013;31(9):814–21. doi: 10.1038/nbt.2676. PubMed PMID: 23975157; PubMed Central PMCID: PMCPMC3819121.

36. Louca S, Doebeli M. Efficient comparative phylogenetics on large trees. Bioinformatics. 2018;34(6):1053–5. Epub 2017/11/02. doi: 10.1093/bioinformatics/btx701. PubMed PMID: 29091997.

37. Parks DH, Tyson GW, Hugenholtz P, Beiko RG. STAMP: statistical analysis of taxonomic and functional profiles. Bioinformatics. 2014;30(21):3123–4. Epub 2014/07/26. doi: 10.1093/bioinformatics/btu494. PubMed PMID: 25061070; PubMed Central PMCID: PMCPMC4609014.

38. White JR, Nagarajan N, Pop M. Statistical methods for detecting differentially abundant features in clinical metagenomic samples. PLoS Comput Biol. 2009;5(4):e1000352. Epub 2009/04/11. doi: 10.1371/journal.pcbi.1000352. PubMed PMID: 19360128; PubMed Central PMCID: PMCPMC2661018.

39. Jeong DY, Yang HJ, Jeong SJ, Kim MG, Yun CY, Lee HY, et al. Immunostimulatory effects of blue-berry yeast fermented powder aginst cyclophosphamide-induced immunosuppressed model. J Physiol & Pathnol Krean Med. 2019;33:48–55.

40. Lee YS, Lee GH, Kwon YK, Park JH, Shin SW. Immunomodulatory effect of aqueous extracted Zingiberis Rhizoma on cyclophosphamide - induced immune suppresion. orean J Oriental Physiology & Pathology. 2007;21:485–90.

41. Lee YS, Lee GH, Park JH, Kwon YK, Shin SW. Water extracted Evodiae Fructus Possesses immunomodulatory activities on cyclophosphamide induced immunesuppression. Korean J Physiology & Pathology. 2007;21:1450–5.

42. Chung JH, Shin PG, Ryu JC, Jang DS, Cho SH. Chemical Compositions of Platycodon grandiflorus (jacquin). A. De Candolle Agric. Chem Biotechnol. 1997;40(2):148–51.

43. Shon MY, Seo JK, Kiom HJ, Sung NJ. Chemical compositions and physiological activities of doraji (Platycodon grandiflorum). J Korean Soc Food SciNutr. 2001;30:717–20.

44. Hong MW. Statistical analyeses of Platycodi Radix prescriptions. Kor J Parmacog. 1974;5:61–7.

45. Kim EH, Gwak JY, Jung MJ. Immunomodulatory activity of Platycodon grandiflorum, Codonopsis lanceolata, and Adenophora triphylla extracts in macrophage cells. J Korean Soc Food Sci Nutr. 2018;47:1069–75.

46. Tohamy AA, El-Ghor AA, El-Nahas SM, Noshy MM. Beta-glucan inhibits the genotoxicity of cyclophosphamide, adriamycin and cisplatin. Mutat Res. 2003;541(1-2):45–53. Epub 2003/10/22. doi: 10.1016/s1383-5718(03)00184-0. PubMed PMID: 14568293.

47. Kirmaz C, Bayrak P, Yilmaz O, Yuksel H. Effects of glucan treatment on the Th1/Th2 balance in patients with allergic rhinitis: a double-blind placebo-controlled study. Eur Cytokine Netw. 2005;16(2):128–34. Epub 2005/06/09. PubMed PMID: 15941684.

48. Naito Y, Uchiyama K, Takagi T. A next-generation beneficial microbe: Akkermansia muciniphila. J Clin Biochem Nutr. 2018;63(1):33–5. Epub 2018/08/09. doi: 10.3164/jcbn.18-57. PubMed PMID: 30087541; PubMed Central PMCID: PMCPMC6064808.

49. Shin NR, Lee JC, Lee HY, Kim MS, Whon TW, Lee MS, et al. An increase in the Akkermansia spp. population induced by metformin treatment improves glucose homeostasis in diet-induced obese mice. Gut. 2014;63(5):727–35. Epub 2013/06/28. doi: 10.1136/gutjnl-2012-303839. PubMed PMID: 23804561.

50. Carbonero F, Benefiel AC, Alizadeh-Ghamsari AH, Gaskins HR. Microbial pathways in colonic sulfur metabolism and links with health and disease. Front Physiol. 2012;3:448. Epub 2012/12/12. doi: 10.3389/fphys.2012.00448. PubMed PMID: 23226130; PubMed Central PMCID: PMCPMC3508456.

51. Kasaikina MV, Kravtsova MA, Lee BC, Seravalli J, Peterson DA, Walter J, et al. Dietary selenium affects host selenoproteome expression by influencing the gut microbiota. FASEB J. 2011;25(7):2492–9. Epub 2011/04/16. doi: 10.1096/fj.11-181990. PubMed PMID: 21493887; PubMed Central PMCID: PMCPMC3114522.

